# Structural Mechanism Underlying Ligand Binding and Activation of PPARγ

**DOI:** 10.1101/2020.09.22.298109

**Authors:** Jinsai Shang, Douglas J. Kojetin

## Abstract

Ligands bind to an occluded orthosteric pocket within the nuclear receptor (NR) ligand-binding domain (LBD). Molecular simulations have revealed several theoretical ligand entry/exit pathways to the orthosteric pocket, but experimentally it remains unclear whether ligand binding proceeds through induced fit or conformational selection mechanisms. Using NMR spectroscopy lineshape analysis, we show that ligand binding to the peroxisome proliferator-activated receptor gamma (PPARγ) LBD involves a two-step induced fit mechanism including an initial fast step followed by slow conformational change. Surface plasmon resonance and isothermal titration calorimetry heat capacity analysis support the fast kinetic binding step and the conformational change after binding step, respectively. The putative initial ligand binding pose is suggested in several crystal structures of PPARγ LBD where a ligand is bound to a surface pore formed by helix 3, the β-sheet, and the Ω-loop—one of several ligand entry sites suggested in previous targeted and unbiased molecular simulations. These findings, when considered with a recent NMR study showing the activation function-2 (AF-2) helix 12 exchanges in and out of the orthosteric pocket in apo/ligand-free PPARγ, suggest an activation mechanism whereby agonist binding occurs through an initial encounter complex with the LBD followed by transition of the ligand into the orthosteric pocket concomitant with a conformational change resulting in a solvent-exposed active helix 12 conformation.

## INTRODUCTION

Nuclear receptors (NRs) comprise a superfamily of transcription factors that evolved to bind and functionally respond to endogenous small molecule ligands (1). NRs contain a conserved domain organization including a central DNA-binding domain flanked by two regulatory regions, a disordered N-terminal activation function-1 (AF-1) domain and a C-terminal ligand-binding domain (LBD) containing the activation function-2 (AF-2) coregulator interaction surface. Endogenous and synthetic ligands bind to an orthosteric pocket within the core of the NR LBD. Ligand binding affects the conformation of the AF-2 surface and changes the binding affinity for chromatin remodeling transcriptional coregulator proteins resulting in activation or repression of gene transcription (2, 3).

Crystal structures have defined static active and inactive/repressive conformations of NR LBDs bound to ligands that enable binding of transcriptional coactivator and corepressor proteins, respectively, by stabilizing specific conformations of the AF-2 helix 12 (4). However, mechanistically it remains poorly understood how ligands engage the LBD and enter the orthosteric ligand-binding pocket—whether ligand binding occurs through conformational selection or induced fit mechanisms (5). In the conformational selection scenario, ligand binding selectively binds to and selects a particular conformation that is populated within the dynamic LBD conformational ensemble. In the induced fit scenario, ligand binding occurs through an encounter complex and induces or pushes the LBD conformational ensemble into the final ligand-bound complex.

In NR LBD crystal structures, ligands bound orthosteric pockets are occluded from solvent suggesting an induced fit binding mechanism. A recent review of molecular simulations on various NR LBDs identified six potential locations involved in ligand entry and exit pathways to the NR orthosteric pocket (6). Most molecular simulation studies focused on ligand egress or unbinding from the orthosteric pocket. However, coarse grained simulations on farnesoid X receptor (FXR) suggest ligand binding occurs through induced fit mechanism (7) via an orthosteric pocket entry site that was also observed in simulations of peroxisome proliferator-activated receptor gamma (PPARγ) (8). Molecular simulations of ligand binding to steroid receptors including androgen receptor (AR), estrogen receptor alpha (ERα), glucocorticoid receptor (GR), mineralocorticoid receptor (MR), and progesterone receptor (PR) also suggest an induced fit mechanism (9, 10), although the site of ligand entry into the orthosteric pocket is different than FXR (7) and PPARγ (8). Indeed, ligand binding to nuclear receptors is often described to induce an active conformation (4, 11–16). However, there is evidence from NMR studies on PPARγ that in the absence of ligand the apo-LBD exchanges between transcriptionally active and transcriptionally inactive/repressive conformations (17) suggesting a role for conformational selection in the ligand binding mechanism of NR agonists, which are thought to stabilize an active conformation from a dynamic ensemble of active and inactive/repressive conformations (15, 18–27). Taken together, these observations stem from the ability of the ligand-bound NR LBD to exert specific functions such as coactivator interaction and transcription, but not directly on the mechanism of ligand binding to the orthosteric pocket.

Here, we use protein NMR lineshape analysis, which provides atomic resolution structural insight into conformational selection and induced fit binding mechanisms (28), to study the mechanism of agonist binding to PPARγ. We find that agonist binding to the PPARγ LBD occurs through an induced fit mechanism with two steps, an initial fast kinetic step followed by a slow conformational change step, which we independently confirm using surface plasmon resonance (SPR) and temperature-dependent isothermal titration calorimetry (ITC) heat capacity analysis, respectively. Crystal structures show that ligands can bind to a surface pore in the PPARγ LBD, a site implicated as a putative ligand entry site in molecular simulations. We discuss the implication of our findings within the context of our recent NMR study showing that helix 12 occupies the orthosteric ligand-binding pocket in apo-PPARγ (17) that when taken together describes a more complete mechanism by which agonist binding induces activation of PPARγ.

## RESULTS

### NMR lineshape analysis of agonist binding

To study the mechanism of ligand binding to PPARγ, we used a previously reported high affinity synthetic PPARγ agonist called GW1929 (**Fig. 1A**) (29) that displays a 100 pM inhibitory binding constant (K_i_) (**Fig. 1B**). We collected 2D [^1^H-^15^N]-TROSY-HSQC NMR spectra of ^15^N-PPARγ LBD in the absence and presence of increasing substoichiometric molar concentrations of GW1929 (**Fig. S1**). For a simple two-state ligand binding mechanism, also called a “lock-and-key” or U model (28), titration of a high affinity ligand with mid-picomolar affinity in principle should result in NMR chemical shift perturbations that occur in slow exchange on the NMR time scale. The intensity of the ligand-free/apo-protein signal would be expected to decrease during the titration while the ligand-bound/holo-protein signal increases. By comparison, titration of a low affinity ligand should cause chemical shift perturbations that occur in fast exchange, causing the apo-protein signal to shift towards the holo-protein signal.

**Fig. 1.**
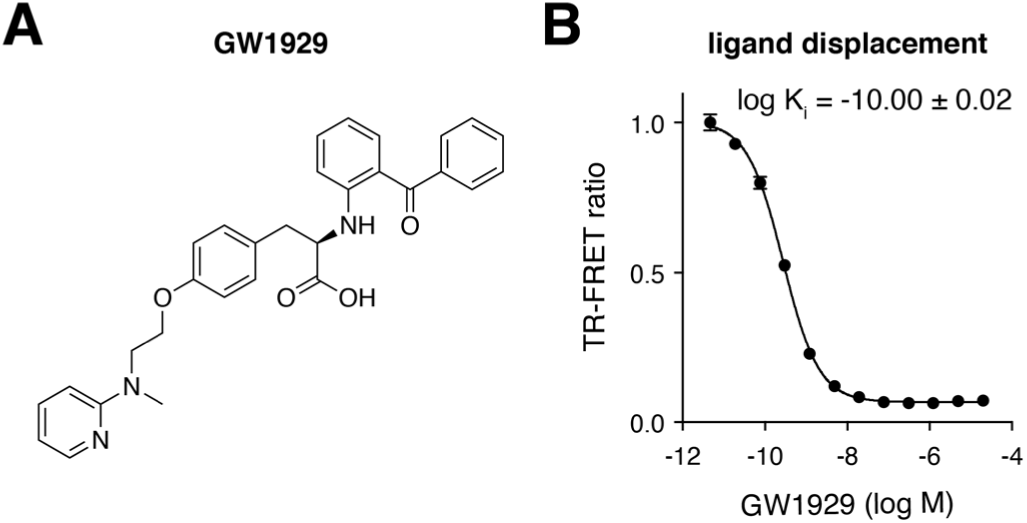
Affinity characterization of GW1929. (**A**) Chemical structure of GW1929. (**B**) TR-FRET fluorescent tracer ligand displacement assay measuring the inhibition constant (K_i_) of GW1929 binding to the PPARγ LBD..

During the GW1929 titration, a mixture of fast and slow exchange NMR lineshapes occur (**Fig. 2A**) for residues dispersed throughout the PPARγ LBD (**Fig. 2B**). The ligand-free apo-peaks disappear in fast exchange, and the ligand-bound holo-peaks are populated in slow exchange. The mixture of fast and slow exchange indicates that GW1929 binding to PPARγ LBD occurs via a three-state mechanism (28) where the protein either isomerizes between two conformations (U-R and U-RL models) or undergoes monomer-dimer equilibrium in the absence or presence of ligand (U-R2 or U-R2L2). Small-angle X-ray scattering (SAXS), dynamic light scattering (DLS), size exclusion chromatography (SEC), and NMR data show that the apo- and ligand-bound PPARγ LBD is monomeric (18, 30), ruling out a potential contribution from protein dimerization (U-R2 and U-R2L2). Furthermore, GW1929 does not contain a racemizable chiral center that would result in an enantiomeric ligand isomerization mixture (U-L) and it is not likely that GW1929 forms a dimer (U-L2), although these features would not contribute to the three-state protein-observed exchange mechanism (28). This leaves two protein isomerization scenarios that show a combination of fast and slow exchange components: ligand binding that occurs via conformational selection or induced fit. In the conformational selection scenario, the protein undergoes slow isomerization and the ligand binds with fast kinetics to a sparsely populated conformation, causing the apo-peak to disappear in slow exchange and the appearance of a holo-peak in fast exchange; this is opposite of what we see in the GW1929 titration data. However, these data are consistent with an induced fit scenario (**Fig. 2C**) where the ligand binds to the protein via a fast kinetic initial step that causes the apo-peak to disappear in fast exchange, which is followed by a slow step where the initial ligand-protein complex changes conformation into a more tightly bound conformation that causes the holo-peak to appear in slow exchange.

**Fig. 2.**
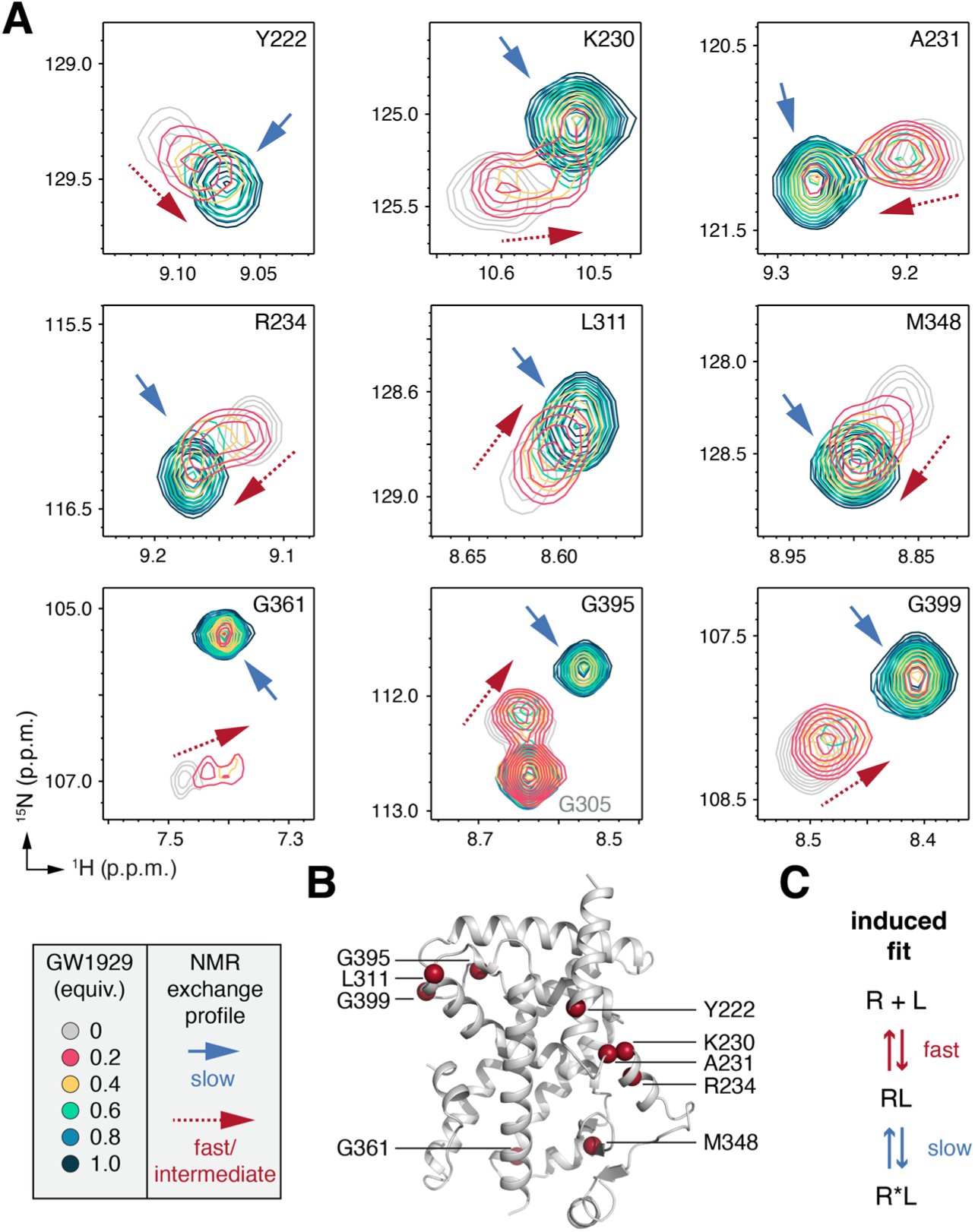
NMR analysis of GW1929 binding to PPARγ LBD. (**A**) Snapshots of 2D [^1^H,^15^N]-TROSY-HSQC NMR spectra of ^15^N-labeled PPARγ LBD in the absence or presence of increasing concentrations of GW1929. (**B**) Structural locations of the residues highlighted in the NMR analysis shown in A. (**C**) A two-state induced fit binding model that explains the mixture of fast and slow exchange lineshapes in the NMR titration data.

### SPR and ITC confirm fast and slow kinetic binding steps

The induced fit binding mechanism suggested by the NMR titration data indicates that the initial ligand binding step involves fast kinetics. To confirm this observation, we performed surface plasmon resonance (SPR) experiments to monitor the binding kinetics of GW1929 to the PPARγ LBD (**Fig. 3A**). As anticipated from the NMR studies, the SPR sensorgram profiles show that GW1929 binds with a fast rate of binding (K_on_ or K_a_). Other published SPR studies of various structurally distinct synthetic PPARγ ligands similarly show fast kinetic rates of binding to the PPARγ LBD (31–42), suggesting that a fast initial binding step may be a common mechanism of ligand binding to the PPARγ.

**Fig. 3.**
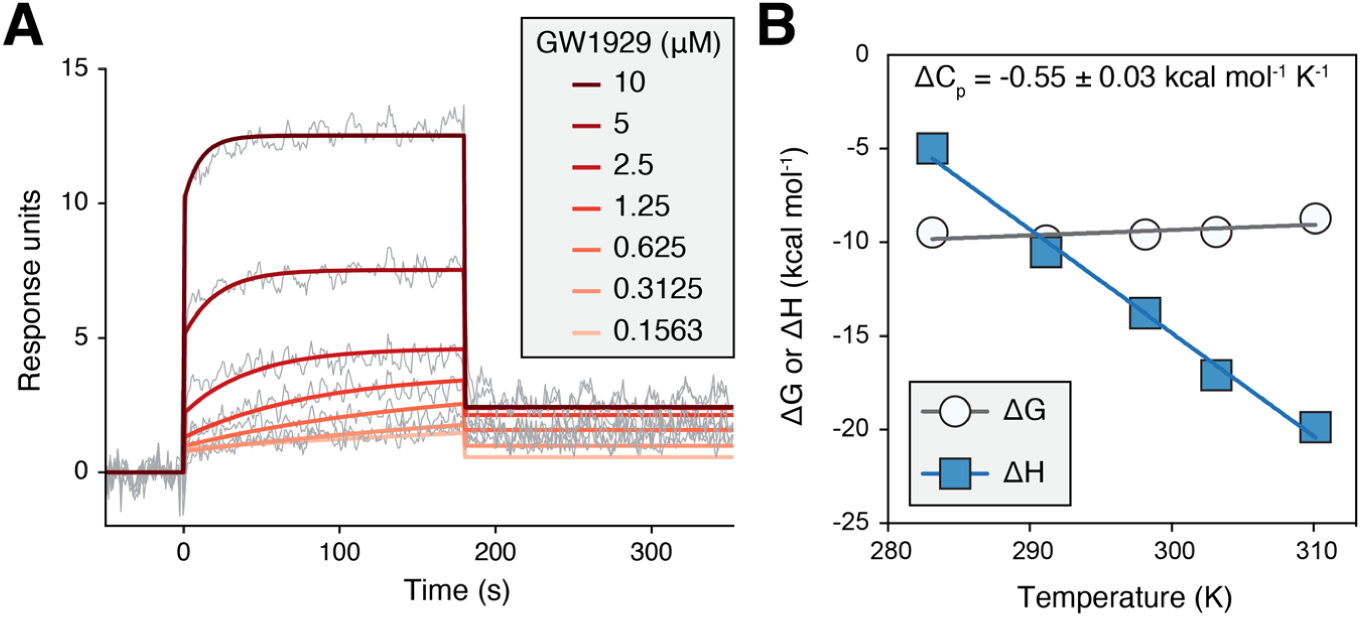
SPR and ITC analysis of GW1929 binding. (**A**) SPR analysis of GW1929 binding to PPARγ LBD shows fast kinetics. (**B**) Heat capacity (ΔC_p_) analysis from temperature-dependent ITC titrations of GW1929 into PPARγ LBD.

To confirm the slow conformational change step that occurs after the initial fast binding step revealed by the NMR studies, we performed isothermal titration calorimetry (ITC) experiments measuring the thermodynamic parameters of GW1929 binding to PPARγ LBD at several temperatures (**Fig. S2**). The magnitude and temperature dependence of the apparent binding heat capacity (ΔC_p_), which is determined from the slope of the apparent binding enthalpy (ΔH) vs. temperature plot, informs whether binding occurs through a lock-and-key (small magnitude, temperature-independent ΔC_p_), conformational selection (larger magnitude, temperature-dependent ΔC_p_), or induced fit (larger magnitude, temperature-independent ΔC_p_) mechanism (43). GW1929 binding to PPARγ LBD shows a strong linear coupling between ΔH vs. temperature (**Fig. 3B**), or a temperature-independent ΔCp, indicative of an induced fit binding mechanism. Taken together, the SPR and ITC data provide support to the NMR data that show GW1929 binds to PPARγ via a two-step induced fit mechanism that includes a fast initial kinetic binding step followed by a slow conformational change.

### Crystal structures reveal the putative ligand entry site to the orthosteric pocket

A common method for obtaining ligand-bound crystal structures of PPARγ LBD is to grow apo-protein crystals and then perform a ligand soak to obtain the ligand-bound complex (44). This procedure is premised on the idea that the path of ligand entry into the orthosteric pocket is accessible to solvent channels within the crystal lattice. To visualize the putative ligand entry site, we grew crystals of apo-PPARγ LBD and solved the structure at 2.27 Å (**Table S1**). Two chains are present in the asymmetric unit (**Fig. S3**) with different helix 12 conformations that are stabilized by a crystal artifact. Helix 12 in the chain A molecule adopts an active conformation and helix 12 in a chain B adopts a non-active conformation because it interacts with the AF-2 surface of a symmetry related chain A molecule. The structure reveals a putative orthosteric pocket entry site: a solvent accessible pore is formed by a surface consisting of helix 3, the β-sheet, and the Ω-loop (**Fig. 4A**). This region was also suggested by molecular dynamics simulations as a putative ligand entry and exit site to the occluded orthosteric ligand-binding pocket in the PPARγ LBD (8, 45).

**Fig. 4.**
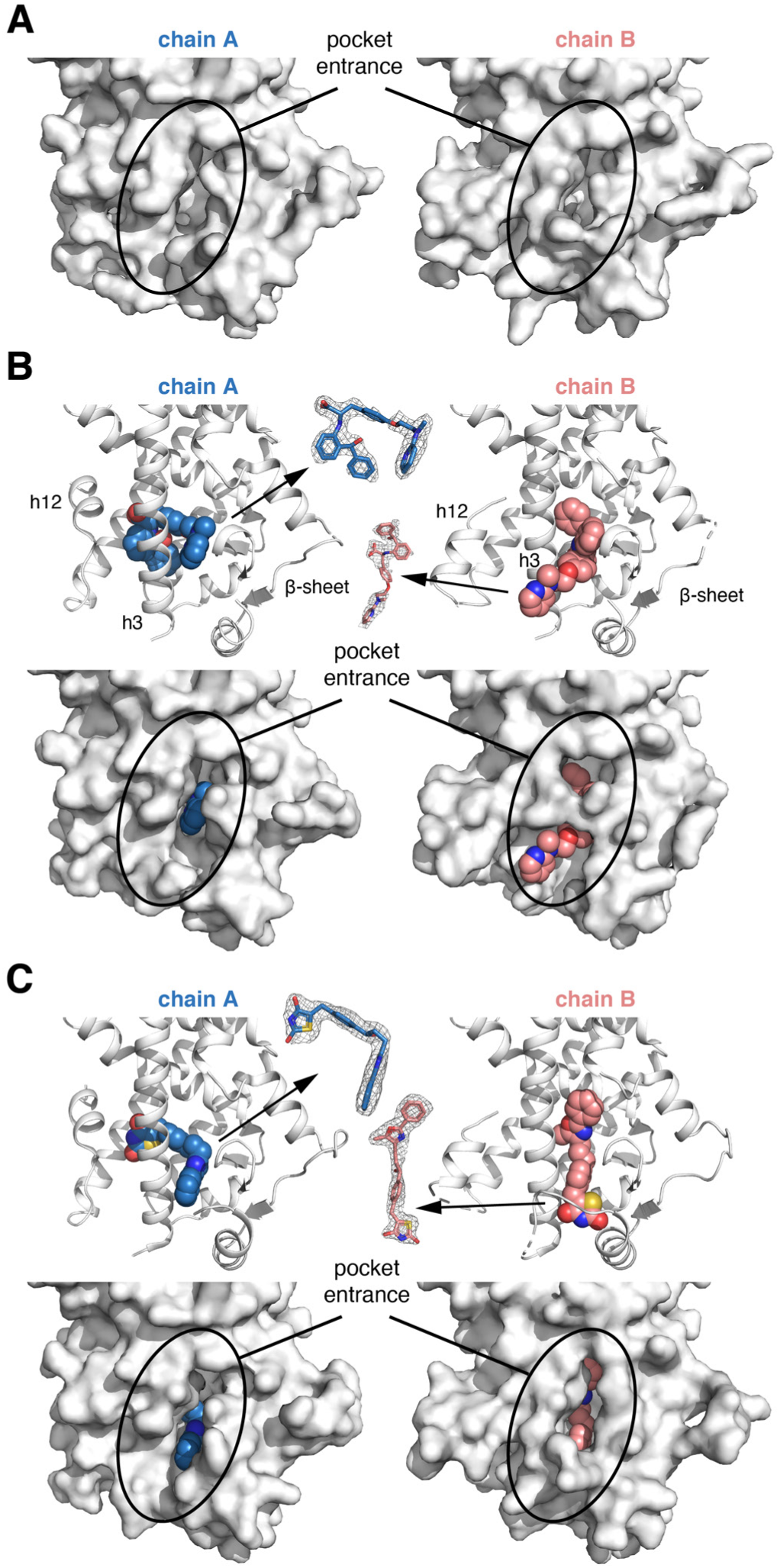
Ligand soaking into apo-PPARγ LBD crystals reveals an orthosteric pocket ligand entry pathway. (**A**) Crystal structure of apo-PPARγ LBD reveals a surface pocket with access to the orthosteric ligand-binding pocket. (**B**,**C**) Soaking of GW1929 (**B**) or darglitazone (**C**) into preformed apo-PPARγ LBD crystals reveals two ligand binding poses, one within the orthosteric pocket in chain A (blue ligand) and a second at the orthosteric pocket ligand entry site (pink ligand). Insets show ligand 2Fo-Fc maps contoured at 1σ, and key structural elements are labeled.

We soaked GW1929 into preformed apo-PPARγ LBD crystals and solved the structure to 2.07 Å (**Table S1**). In chain A where helix 12 is stabilized in an active conformation, GW1929 adopts a binding mode within the orthosteric ligand-binding pocket as typical of PPARγ agonists (**Fig. 4B**). However, in chain B where helix 12 adopts a crystal contact-induced non-active helix 12 conformation, GW1929 bound to the putative orthosteric pocket entry site. Other ligand-bound PPARγ LBD crystal structures obtained from soaking ligand into apo-protein crystals have similarly revealed a ligand bound to this pocket entrance in chain B molecules (**Fig. S4**) (24, 37, 46–51), including a crystal structure we previously solved of darglitazone-bound PPARγ LBD (**Fig. 4C**). These crystallography observations provide support that this solvent accessible pore at the helix ^I^ 3/β-sheet/Ω-loop surface constitutes the ligand entry site to the PPARγ orthosteric pocket.

### Induced fit as a general PPARγ ligand binding mechanism

The data above raise a question as to whether other structurally distinct synthetic PPARγ agonists bind using an induced fit mechanism. To address this we used NMR to study the binding mechanism of a different full agonist, darglitazone, and a partial agonist, MRL24 (**Fig. 5A**), which display K_i_ values at or below 1 nM in a ligand displacement assay (**Fig. 5B**). We collected 2D [^1^H-^15^N]-TROSY-HSQC NMR spectra of ^15^N-PPARγ LBD in the absence and presence of substoichiometric molar concentrations of the high affinity agonists. Similar to the NMR titration of GW1929, we observed a mixture of fast and slow exchange NMR lineshapes upon titration of darglitazone (**Fig. 5C, Fig. S5**) and MRL24 (**Fig. 5C, Fig. S6**) associated with the disappearance of ligand-free apo-peaks (fast exchange) and appearance of ligand-bound holo-peaks (slow exchange). Taken together with the GW1929 studies, these data indicate that the induced fit binding mechanism may be a general ligand binding mechanism that is not an artifact associated with any one specific PPARγ ligand scaffold.

**Fig. 5.**
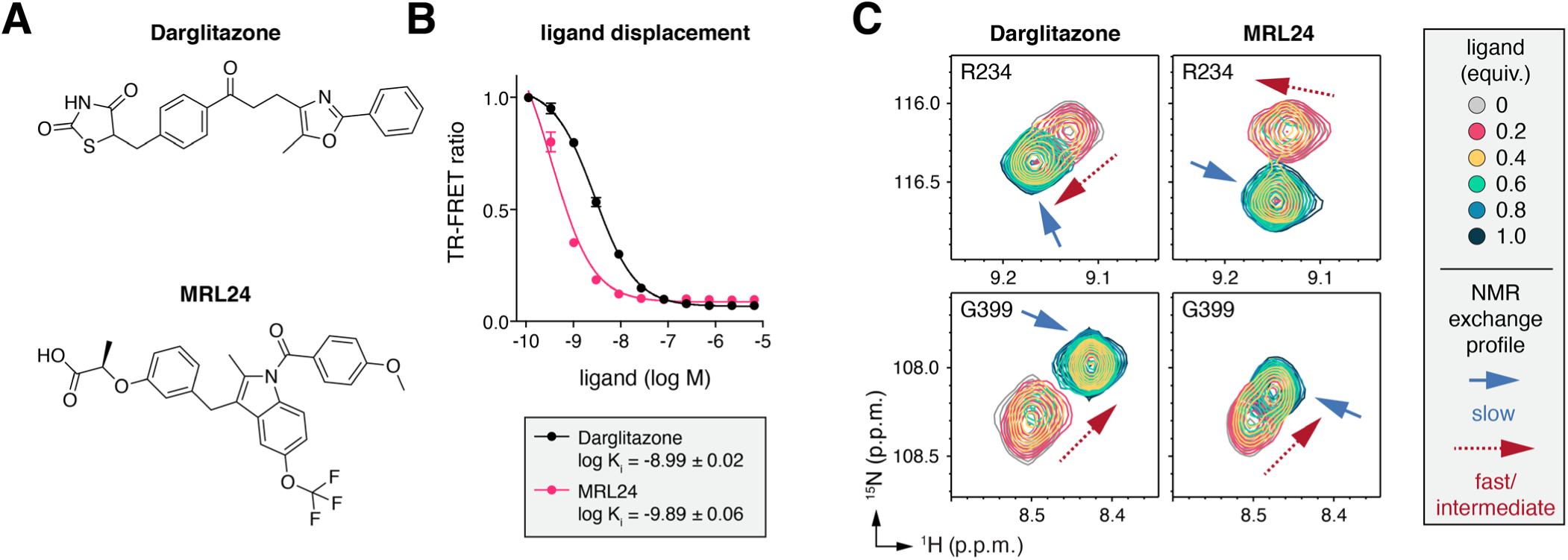
NMR analysis of two structurally distinct ligands binding to PPARγ LBD. (**A**) Chemical structures of darglitazone and MRL24. (**B**) TR-FRET fluorescent tracer ligand displacement assay measuring the inhibition constant (K_i_) of darglitazone and MRL24 binding to the PPARγ LBD. (**C**) Snapshots of 2D [^1^H,^15^N]-TROSY-HSQC NMR spectra of ^15^N-labeled PPARγ LBD in the absence or presence of increasing concentrations of darglitazone or MRL24.

### A helix 12 mutant impairs full agonist-induced function but not binding mechanism

When bound to the orthosteric pocket, full agonists such as darglitazone and GW1929 form a hydrogen bond with the side chain hydroxyl group of residue Y473 on helix 12. This tyrosine residue is thought to be critical for binding affinity and transcriptional efficacy of full agonists, but not partial agonists that do not interact with Y473 and display lower levels of PPARγ-mediated transcription (52). The crystal structures of wild-type PPARγ LBD with the non-active helix 12 conformation observed in chain B provide one glimpse into a ligand encounter complex when helix 12 does not adopt an active conformation that points the Y473 side chain into the orthosteric pocket. To further determine how Y473 impacts the binding mechanism of darglitazone and GW1929, we generated a mutant [Y473E]-PPARγ LBD construct that we hypothesized would impact the ligand binding model in both chains A and B. In a TR-FRET coregulator interaction assay, the Y473E mutation significantly decreases the darglitazone EC_50_ and efficacy for increasing interaction of the TRAP220/MED1 coactivator peptide, and GW1929 shows essentially no concentration-dependent effect on the interaction (**Fig. 6A**).

**Fig. 6.**
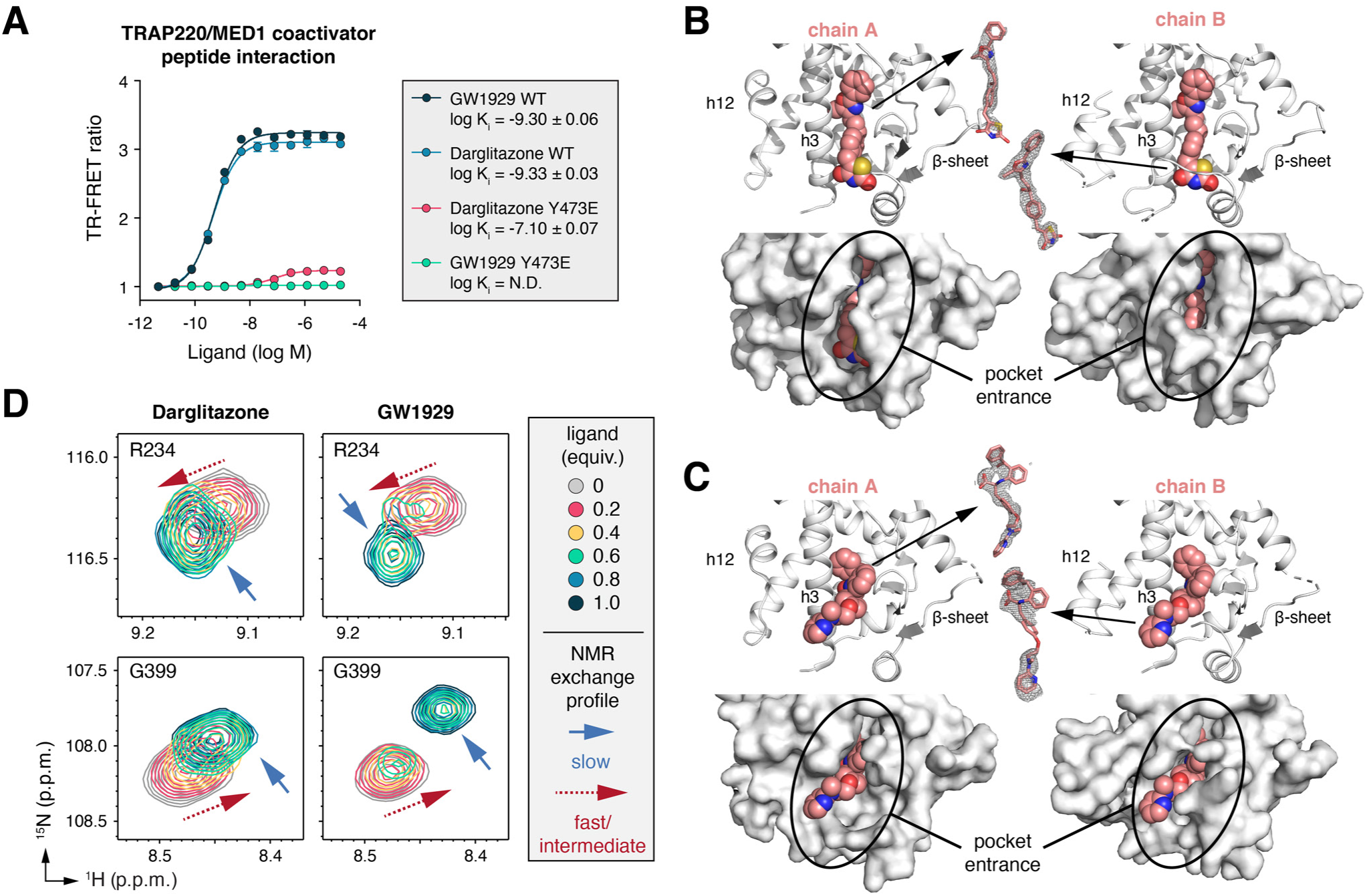
Ligand binding analysis to [Y473E]-PPARγ LBD. (**A**) TR-FRET coregulator interaction assay characterizing the activity of two full agonists, darglitazone and GW1929, on the interaction of a peptide derived from the TRAP220/MED1 coactivator protein to either wild-type (WT) PPARγ LBD or [Y473E]-PPARγ LBD. (**B**,**C**) Soaking of darglitazone (**B**) or GW1929 (**C**) into preformed apo-[Y473E]-PPARγ LBD crystals reveals the ligands bind only to the orthosteric pocket ligand entry site in both chain A and B (pink ligand). Insets show ligand 2Fo-Fc maps contoured at 1σ, and key structural elements are labeled.

We crystallized [Y473E]-PPARγ LBD under the same conditions used to generate apo-PPARγ LBD crystals and solved the structure at 2.30 Å (**Table S1**). Overall, the [Y473E]-PPARγ LBD structure is highly similar to the apo-PPARγ LBD structure; chain A and B in both structures adopt an active and inactive conformation with Cα R.M.S.D values of 0.36 Å and 0.40 Å to the wild-type chain A and B conformers, respectively. We also solved structures after soaking darglitazone (**Fig. 6B**) and GW1929 (**Fig. 6C**) into preformed [Y473E]-PPARγ LBD crystals to 2.40 Å and 2.15 Å (**Table S1**), respectively. Ligand density is present at the ligand entry site but not in the orthosteric pocket in both chain A and B, indicating the Y473 side chain may be necessary for transition of the ligands into the final high affinity orthosteric binding pose within the crystals.

Although the TR-FRET and crystallography data indicate that the Y473E mutant inhibits or weakens GW1929 and darglitazone binding to the orthosteric pocket, 2D [^1^H-^15^N]-TROSY-HSQC NMR spectra of ^15^N-[Y473E]-PPARγ LBD titrated with substoichiometric molar concentrations of darglitazone (**Fig. S7**) and GW1929 (**Fig. S8**) reveals the ligands indeed bind to [Y473E]-PPARγ LBD. Moreover, NMR lineshape analysis of the titration series shows a mixture of fast and slow exchange indicating that binding of darglitazone and GW1929 (**Fig. 6D**) proceeds through an induced fit mechanism to [Y473E]-PPARγ LBD similar to the binding mechanism to wild-type PPARγ LBD. The slow exchange characteristic of the NMR titration profile suggests that the ligands bind to [Y473E]-PPARγ LBD with a reasonably high affinity, which we confirmed for GW1929 using ITC (**Fig. S9**). Overlay of NMR spectra of wild-type or Y473E mutant ^15^N-PPARγ LBD bound to 1 equivalent of darglitazone (**Fig. S10**) or GW1929 (**Fig. S11**) look very similar with two notable exceptions. First, NMR peaks corresponding to residues in the AF-2 surface, in particular helix 12 but also helix 3–5 and other nearby structural elements, are present in wild-type spectra but missing in the Y473E mutant spectra due to dynamics on the µs-ms intermediate exchange NMR time scale. Second, NMR peaks corresponding to residues within or nearby the AF-2 surface display chemical shift perturbations, likely due to the fact that helix 12 is dynamic and samples multiple conformations in the Y473E mutant but in wild-type PPARγ LBD it is stabilized in an active conformation via hydrogen bond formation between the ligand and the Y473 hydroxyl group.

Taken together, the Y473E ligand binding data are consistent with a dynamic activation model (3) whereby PPARγ agonism is associated with ligand-induced stabilization of helix 12, which is dynamic on the µs-ms time scale in apo-PPARγ (18, 23–25). The NMR data indicate that darglitazone and GW1929 binding to [Y473E]-PPARγ LBD stabilizes the dynamics of most of the orthosteric ligand-binding pocket, likely by binding to the orthosteric pocket in solution though not in the crystallized form. However, because the ligands do not hydrogen bond to the side chain of Y473E, helix 12 remains dynamic, which could explain the relatively flat ligand dose response curve in the TR-FRET assay despite their ability to bind with relatively high affinity. It is also possible that the addition of a coactivator peptide forces the [Y473E]-PPARγ LBD into an active AF-2 helix 12 conformation resulting in a clash between the glutamic acid side chain and the acid headgroup of GW1929 in the orthosteric binding pose. An electrostatic clash in the [Y473E]-PPARγ LBD crystals, or lack of the Y473 side chain in chain B of wild-type PPARγ crystals, could also explain why the ligands do not crystallize with an orthosteric binding pose. However, the NMR and ITC data indicate that the Y473E mutant does not prevent the ligands from binding to the orthosteric pocket in solution with high affinity.

## DISCUSSION

There is evidence that the functional activity of ligand-bound NRs is associated with a shift in the dynamic LBD conformational ensemble from a ground state to an active state. For example, in the absence of ligand, the PPARγ LBD is conformationally dynamic, samples multiple conformation, and binding of an agonist stabilizes the LBD in an active conformation to a degree correlated with the activity of the ligand (18, 23–25). The association between ligand-bound NR LBD conformation and graded function is evidence for conformational selection in the mechanism of NR ligand activity (13, 15, 19–22, 53). However, these studies only address the functional mechanism of the ligand-bound state; they do not address the mechanism of ligand binding. In this study, we directly probed the mechanism of ligand binding to PPARγ using NMR lineshape analysis, a powerful method for studying binding equilibria at atomic resolution that can differentiate conformational selection and induced fit binding mechanisms (28).

Our NMR data show that agonists bind to the PPARγ orthosteric pocket via a two-state induced fit mechanism. The first step is characterized by a fast binding event, which is supported by our SPR data and likely occurs at a surface pore formed by helix 3, the β-sheet, and the Ω-loop. This is a site where ligands can bind in the non-active conformation PPARγ LBD crystal structures as well as our Y473E mutant crystal structures, and it is the ligand entry site suggested in unbiased coarse-grained molecular simulations of FXR (7) and targeted simulations of PPARγ (8). Molecular simulations of other NRs have suggested other ligand entry sites including a conserved surface in steroid receptors at the intersection of helix 3, 7, and 11 (9, 10), suggesting that certain classes of NRs may use different ligand entry sites to the orthosteric pocket.

The second slower step detected by our NMR studies is associated with a conformational change after the initial fast binding event, which is supported by our ITC heat capacity analysis. This step likely represents transition of the ligand from the initial encounter complex at the surface pore ligand entry site to the final bound conformation within the orthosteric pocket. Molecular simulation studies of ligand binding to FXR (7) and PPARγ (8) support this conformational change step. In the study on PPARγ, binding of an agonist called GW0072 was described to rotate on itself during a transition from its initial binding pose to the final bound conformation. This is conceptually similar to the crystallized PPARγ ligand binding modes we describe here. In the structures of darglitazone bound to the pocket entry site, the TZD headgroup is solvent exposed—if this represents an initial encounter complex binding pose that is populated in solution, the TZD headgroup would need to swing around helix 3 through a rotation point at the intersection of helix 3 and 5 to migrate to the final orthosteric binding pose. In the structures of GW1929 bound to the pocket entry site, the acid headgroup points into the pocket and if this also represents an initial encounter complex binding pose that is populated in solution it would need to flip ∼180° during the transition into the final orthosteric binding pose. The unbiased coarse-grained simulations of obeticholic acid binding to FXR revealed a similar multi-step mechanism with the final step associated with a rearrangement of the FXR LBD and a transition of the ligand to the inner binding pose within the orthosteric pocket (7).

Our findings extend a model for the molecular mechanism of agonist binding and activation of PPARγ (**Fig. 7**). In the absence of ligand, several regions of apo-PPARγ LBD exchanges between multiple conformations on the microsecond-millisecond (µs-ms) time scale including the orthosteric ligand-binding pocket and the AF-2 helix 12 (18). One of these conformations was assumed to be the active NR LBD conformation that has been captured in most NR crystal structures. Using paramagnetic relaxation enhancement (PRE) NMR, we recently showed for apo-PPARγ LBD that the µs-ms time scale dynamics is caused by exchange of the AF-2 helix 12 in and out of the orthosteric ligand-binding pocket, which are associated with the transcriptionally repressive and active conformations (17). This suggests that there may be a competition between the repressive helix 12 conformation within the orthosteric pocket and the transition of the ligand from the encounter complex bound at the entry site to the final binding pose in the orthosteric pocket. It is possible that this transition pushes helix 12 out of the repressive conformation within the orthosteric pocket to a solvent-exposed active conformation, a signature of what might be considered a true induced fit binding mechanism. Alternatively, the binding mechanism could involve a mixture of induced fit binding and conformational selection where helix 12 may adopt an active-like conformation prior to the ligand transitioning to the final binding pose in the orthsoteric pocket if the time scale by which helix 12 exchanges out of the pocket is not rate limiting (5). Other NMR studies have similarly found that the orthosteric pocket and helix 12 in other apo-NR LBDs are dynamic on the µs-ms time scale including FXR, PPARα, retinoid X receptor alpha (RXRα), and vitamin D receptor (VDR) (19–21, 27). Thus, the ligand binding mechanism we describe here for PPARγ may occur for other NRs as well, a hypothesis that could be tested by NMR and biophysical studies as demonstrated in this study.

**Fig. 7.**
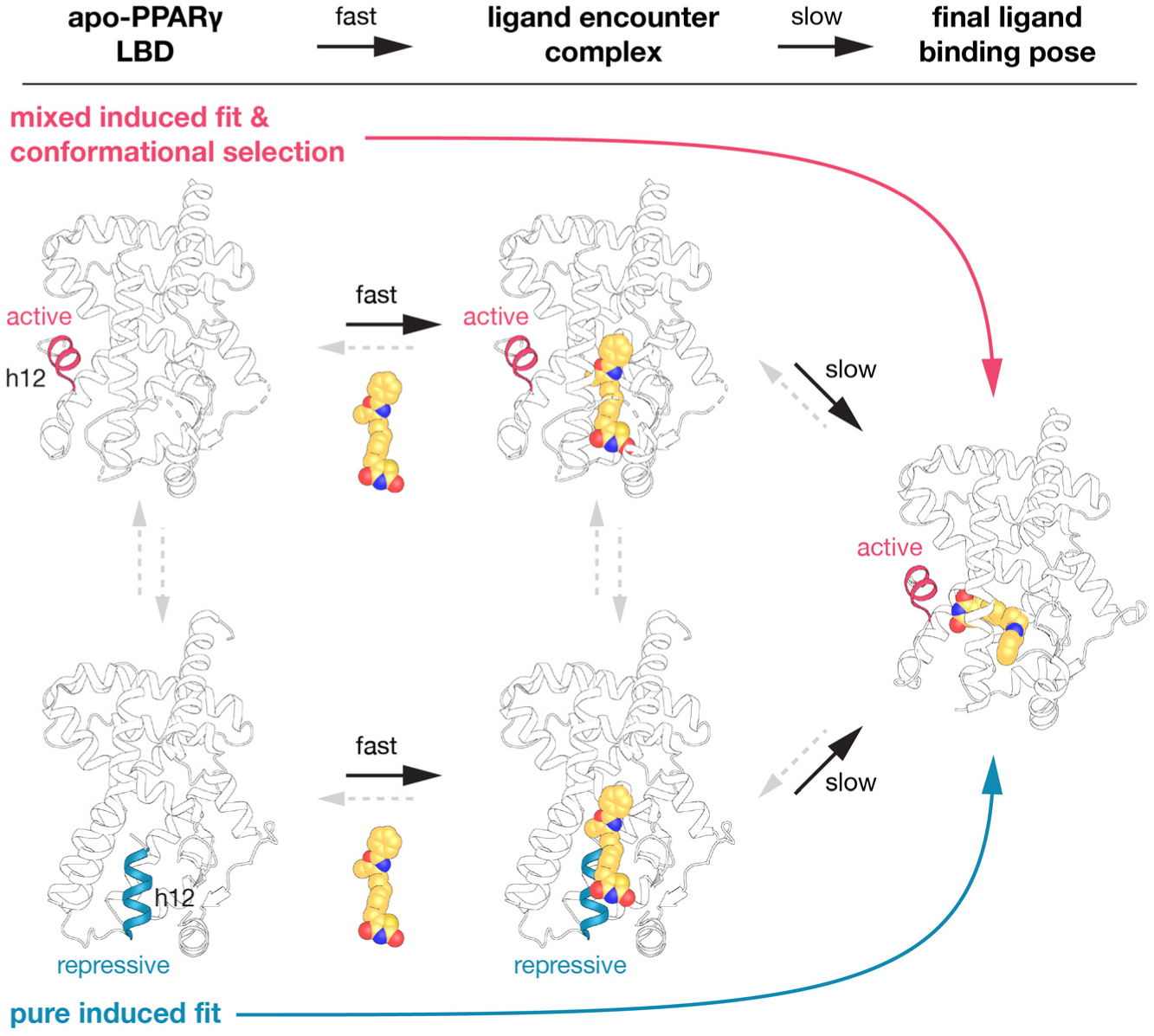
Schmatic model of agonist binding to a dynamic apo-PPARγ LBD conformational ensemble via a two-step process involving an initial fast step followed by a slow conformational change step. In the absence of ligand, helix 12 in apo-PPARγ LBD exchanges between a transcriptionally repressive conformation within the orthosteric ligand-binding pocket and a solvent exposed active conformation. Agonist binding to the ligand entry site via an encounter complex could occur to either of these conformations. In the pure induced fit scenario, agonist binding to the repressive LBD conformation would push helix 12 into an active conformation. In the mixed induced fit & conformational selection scenario, ligand binding to the active LBD conformation would facilitiate transition into the final ligand binding pose. Structures displayed in this schmatic model include PDB code 6ONJ (active conformation; chain A), 6ONI (repressive conformation; chain B), and 6DGL (darglitazone ligand; chain B). Black arrows denote processes supported by experimental studies (NMR, SPR, and ITC heat capacity analysis); grey arrows denote other steps that are part of the binding kinetic pathways.

## ACKNOWLEDGEMENTS

This work was supported by National Institutes of Health (NIH) grant R01DK124870. Use of the Stanford Synchrotron Radiation Lightsource, SLAC National Accelerator Laboratory, is supported by the U.S. Department of Energy, Office of Science, Office of Basic Energy Sciences under Contract No. DE-AC02-76SF00515. The SSRL Structural Molecular Biology Program is supported by the DOE Office of Biological and Environmental Research, and by the National Institutes of Health, National Institute of General Medical Sciences (including P41GM103393). The contents of this publication are solely the responsibility of the authors and do not necessarily represent the official views of NIDDK or NIH.

## AUTHOR CONTRIBUTIONS

J.S. and D.J.K. conceived and designed the research. J.S. performed all experiments and analyzed data. D.J.K. analyzed data and supervised the research. J.S. and D.J.K. wrote the manuscript.

## METHODS

### Materials and reagents

Darglitazone and GW1929 were obtained from Cayman Chemical, MRL24 was obtained from BioVision; ligands were dissolved in DMSO-d_6_ at 20 mM stocks. Peptides of LXXLL-containing motif from human TRAP220/ MED1 (UniProt Q15648; residues 638–656; NTKNHPMLMNLLKDNPAQD) with an N-terminal FITC label, a six-carbon linker (Ahx), and an amidated C-terminus for stability was synthesized by LifeTein.

### Protein expression and purification

Wild-type or Y473E mutant human PPARγ LBD (UniProt P37231; residues 203–477, isoform 1 numbering) protein was expressed from a pET46 Ek/LIC vector (Novagen) as a Tobacco Etch Virus (TEV)-cleavable N-terminal Hexa(6x)His-tagged fusion protein in *Escherichia coli* BL21(DE3) cells using autoinduction ZY media (grown at 22°C; harvested after overnight growth), or M9 minimal media supplemented with ^15^NH_4_Cl (induced with 0.4mM isopropyl β-d-1-thiogalactopyranoside at O.D. ∼0.8; grown at 18°C; harvested after overnight growth). The Y473E PPARγ LBD mutant was generated with site directed mutagenesis using the following forward primer and the corresponding reverse primer: 5’-CTGCAGGAGATCGAAAAGGACTTGTAC-3’. Proteins were purified using Ni-NTA affinity chromatography, in some cases a second Ni-NTA affinity chromatography step after cleavage of the 6xHis-tag by TEV protease (at a ratio of 1 mg TEV protease: 50 mg of PPARγ LBD), and finally gel filtration chromatography. Purified protein was concentrated to 10 mg/mL in a buffer consisting of 20 mM potassium phosphate (pH 7.4), 50 mM potassium chloride, 5 mM tris(2-carboxyethyl)phosphine (TCEP), and 0.5 mM ethylenediaminetetraacetic acid (EDTA). Purified proteins were verified by SDS-PAGE as >95% pure. Delipidated protein was obtained by denaturation using a chloroform/methanol lipid extraction method and refolding using a fast dilution/dialysis procedure followed by size exclusion chromatography (54).

### TR-FRET assays

The time-resolved fluorescence resonance energy transfer (TR-FRET) assays were performed in black 384-well plates (Greiner). Ligand stocks were prepared via serial dilution in DMSO and added to wells. For the TR-FRET ligand displacement assay, each well (23 µL total volume) contained 1 nM 6xHis-tagged PPARγ LBD protein, 1 nM LanthaScreen Elite Tb-anti-HIS antibody (Thermo Fisher), and 5 nM Fluormone Pan-PPAR Green fluorescent tracer ligand (Invitrogen) in the same buffer. For the coregulator interaction assay, each well (23 µL total volume) contained 4 nM 6xHis-tagged PPARγ LBD, 1 nM LanthaScreen Elite Tb-anti-HIS antibody (Thermo Fisher), and 400 nM FITC-labeled TRAP220 or NCoR peptide in a buffer consisting of 20mM potassium phosphate (pH 8), 50 mM potassium chloride, 5 mM TCEP, and 0.005% Tween 20. Plates were incubated at 25°C for 1 h and read using Synergy Neo plate reader (BioTek). The Tb donor was excited at 340 nm, its emission was monitored at 492 nm, and the acceptor FITC emission was measured at 520 nm. Data were analyzed using GraphPad Prism. TR-FRET coregulator data were fit to sigmoidal dose response curve equation, and ligand displacement data were fit to the “one site – Fit Ki” binding equation to obtain K_i_ values using the published binding affinity of Fluormone Pan-PPAR Green (2.8 nM; Invitrogen PV4894 info sheet).

### NMR spectroscopy

Two dimensional (2D) [^1^H,^15^N]-transverse relaxation optimized spectroscopy (TROSY) heteronuclear single quantum coherence (HSQC) data were collected on a Bruker 700 MHz NMR spectrometer equipped with a QCI cryoprobe at 298K using Topspin 3.0 (Bruker). Samples contained 200 µM ^15^N-PPARγ LBD at increasing ligand concentrations (0, 0.2, 0.4, 0.6, 0.8, and 1.0 equivalents); a DMSO control experiment (1.0 equivalent of vehicle) showed no significant perturbations. Data were processed using NMRFx Processor (55) and analyzed using NMRViewJ and NMRFx Analyst (56). Backbone NMR chemical shift assignments for GW1929-bound PPARγ LBD were previously reported (57).

### Surface plasmon resonance

Surface plasmon resonance (SPR) measurements were performed on a Biacore X100 instrument (GE Healthcare). 6xHis-tagged PPARγ LBD protein was immobilized on an NTA sensor chip (GE Healthcare), which includes a reference flow cell for background subtraction, at a density not exceeding 2,000 RU using reagents supplied by NTA reagent kit (GE Healthcare). Measurements were performed in a buffer (1X HBS-P+ buffer) containing 10 mM HEPES, 150 mM NaCl and 0.05% (v/v) Surfactant P20 at a flow rate of 10 μL/min. For kinetic and affinity measurements, GW1929 was diluted in 1X HBS-P+ buffer and injected at 8 concentrations with one duplicate. The NTA sensor chip was regenerated between each GW1929 concentration measurement by stripping Ni^2+^ using EDTA and recharging with NiCl_2_.

### Isothermal titration calorimetry

Isothermal titration calorimetry (ITC) experiments were carried out on a iTC200 calorimeter (MicroCal/GE/Malvern) using the iTC200 software (v 1.24.2) for instrument control and data acquisition. GW1929 (present in the syringe at 300 µM) was diluted in a buffer consisting of 20 mM potassium phosphate (pH 7.4), 50 mM potassium chloride, 5 mM TCEP, 0.5 mM EDTA, and 0.17% DMSO. Wild-type or Y473E PPARγ LBD protein (present in the sample cell at 30 µM) was diluted in the same buffer. Titrations were performed with a 0.4 μL pre-injection followed by nineteen 2 μL injections. Mixing was carried out at five temperatures (10°C, 18°C, 25°C, 30°C, 37°C) for wild-type PPARγ LBD or 25°C for [Y473E]-PPARγ LBD with reference power and rotational stirring set at 5 μcal s^-1^ and 1200 rpm, respectively. Data analysis was performed using software packages NITPIC, SEDPHAT, and GUSSI (58–61).

### Crystallization, structure determination, and structural analysis

Crystals of wild-type and Y473E PPARγ LBD were obtained after 3–8 days at 22°C by sitting-drop vapor diffusion against 50 μL of well solution using 96-well format crystallization plates. The crystallization drops contained 1 μL of protein/ligand sample mixed with 1 μL of reservoir solution containing 0.1 M MOPS (pH 7.6) and 0.8 M sodium citrate. Crystals for the darglitazone and GW1929-PPARγ LBD complexes were obtained by soaking ligands (1.5 mM in reservoir solution containing 5% DMSO) into preformed apo-PPARγ LBD crystals for about 1 week. All crystals were flash-frozen in liquid nitrogen before data collection. Data collection for the wild-type apo-refolded and GW1929-bound structures were collected at Berkeley Center for Structural Biology beamline 5.0.3; the apo-Y473E mutant and GW1929 or darglitazone-bound Y473E mutant structures were collected at SLAC National Accelerator Laboratory/Stanford Synchrotron Radiation Lightsource (SSRL) beamline 12-2. Data were processed, integrated, and scaled with the programs Mosflm (62) and Scala in CCP4 (63). The structure was solved by molecular replacement using the program Phaser (64) that is implemented in the PHENIX package (65) using previously published PPARγ LBD structure (PDB code 1PRG) (66) as the search model. The structure was refined using PHENIX with several cycles of interactive model rebuilding in Coot (67). The align command in PyMOL (Schrödinger) was used to calculate Cα R.M.S.D. values with the number of alignment refinement cycles set to 0.

