## supplementary information for "Structural Mechanism Underlying Ligand Binding and Activation of PPARγ"

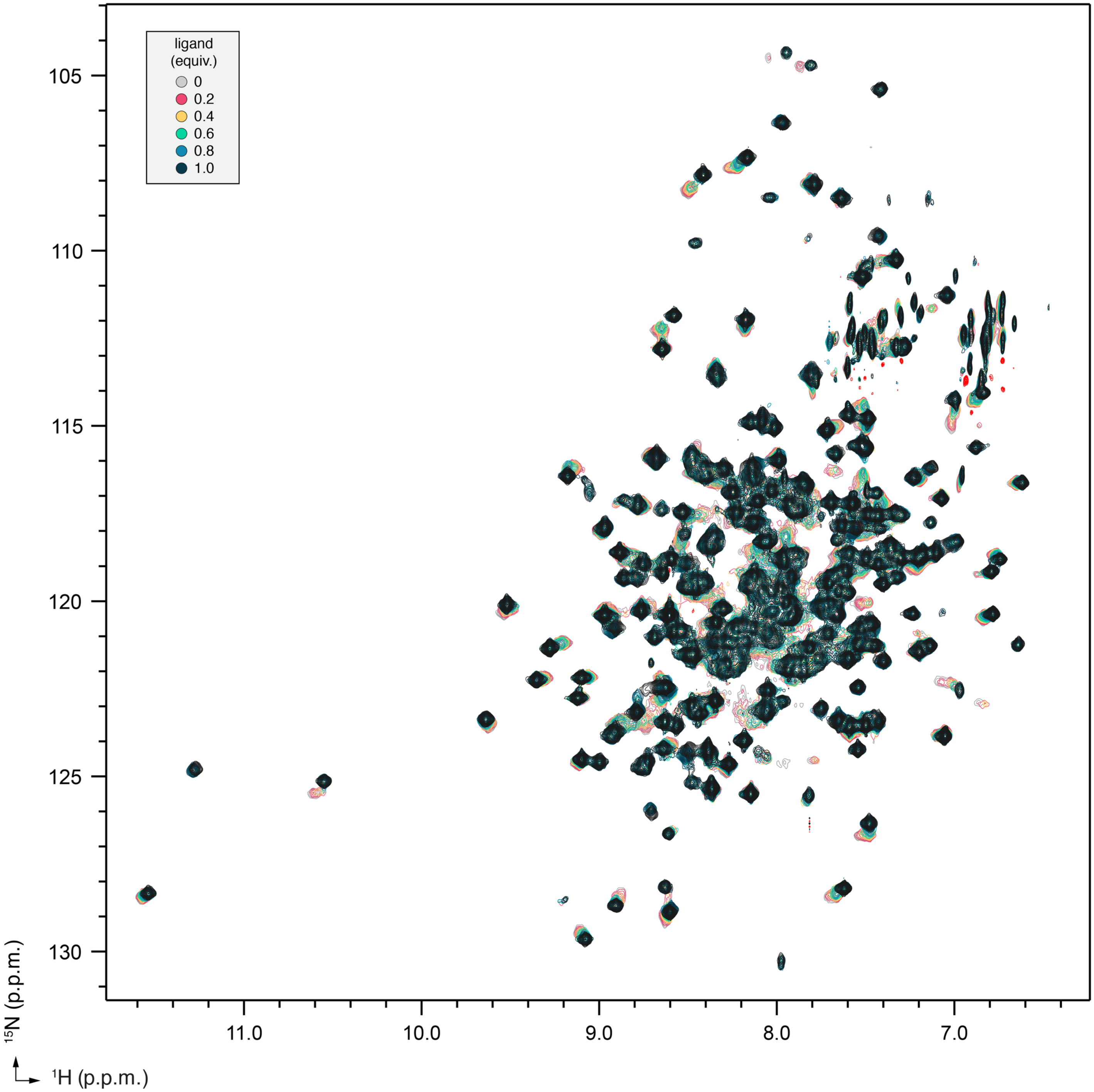

**Fig. S1.** 2D [$^1$H,$^{15}$N]-TROSY-HSQC NMR data of GW1929 titrated into $^{15}$N-PPARγ LBD at the indicated molar equivalents.

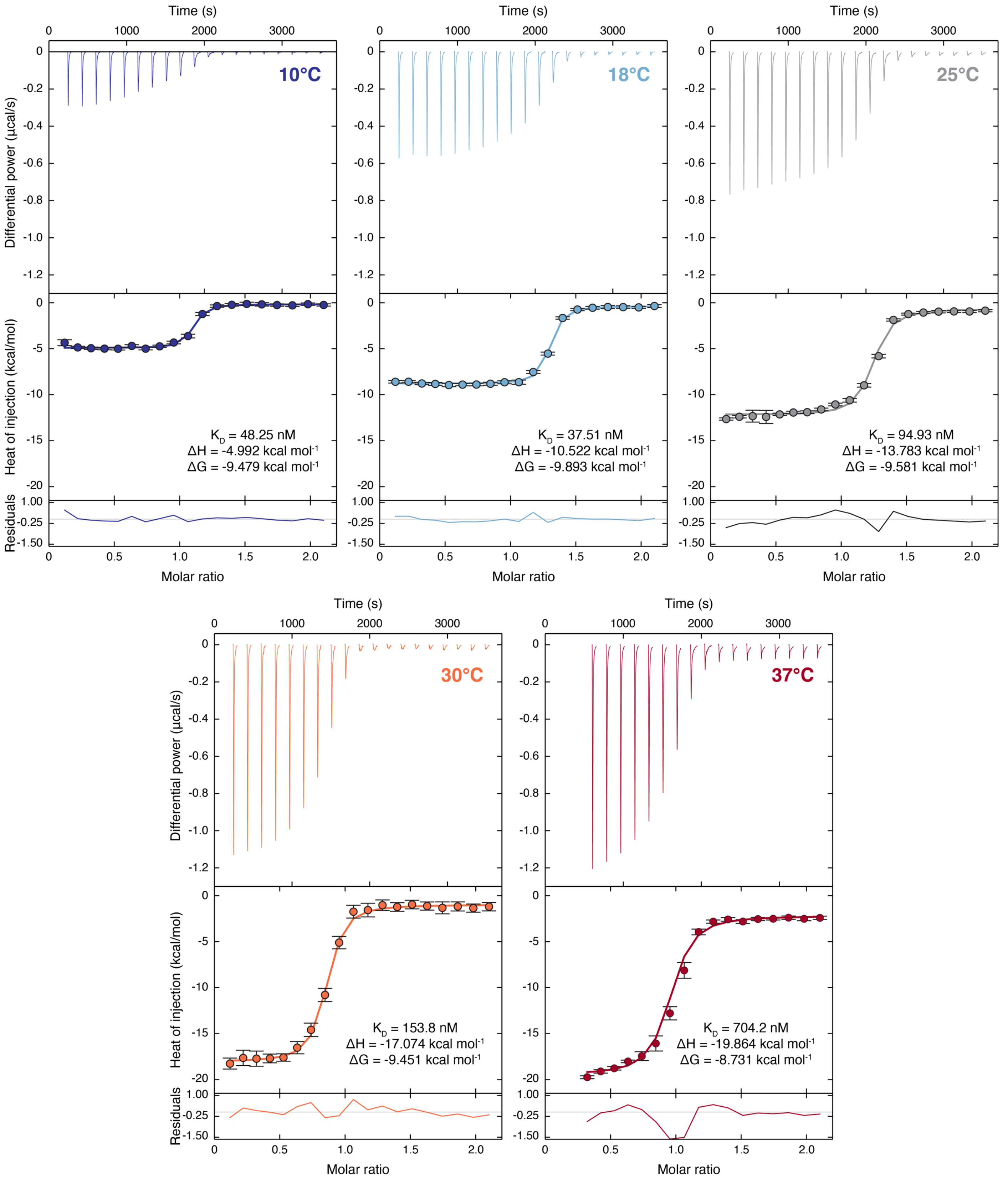

**Fig. S2.** ITC data of GW1929 titrated into PPARγ LBD at the indicated temperatures.

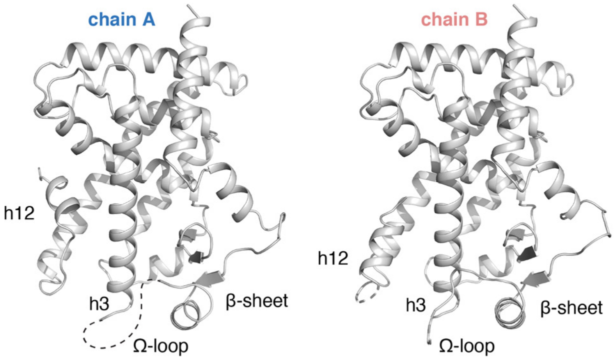

**Fig. S3.** Cartoon diagrams of apo-PPARγ LBD chain A and B molecules.

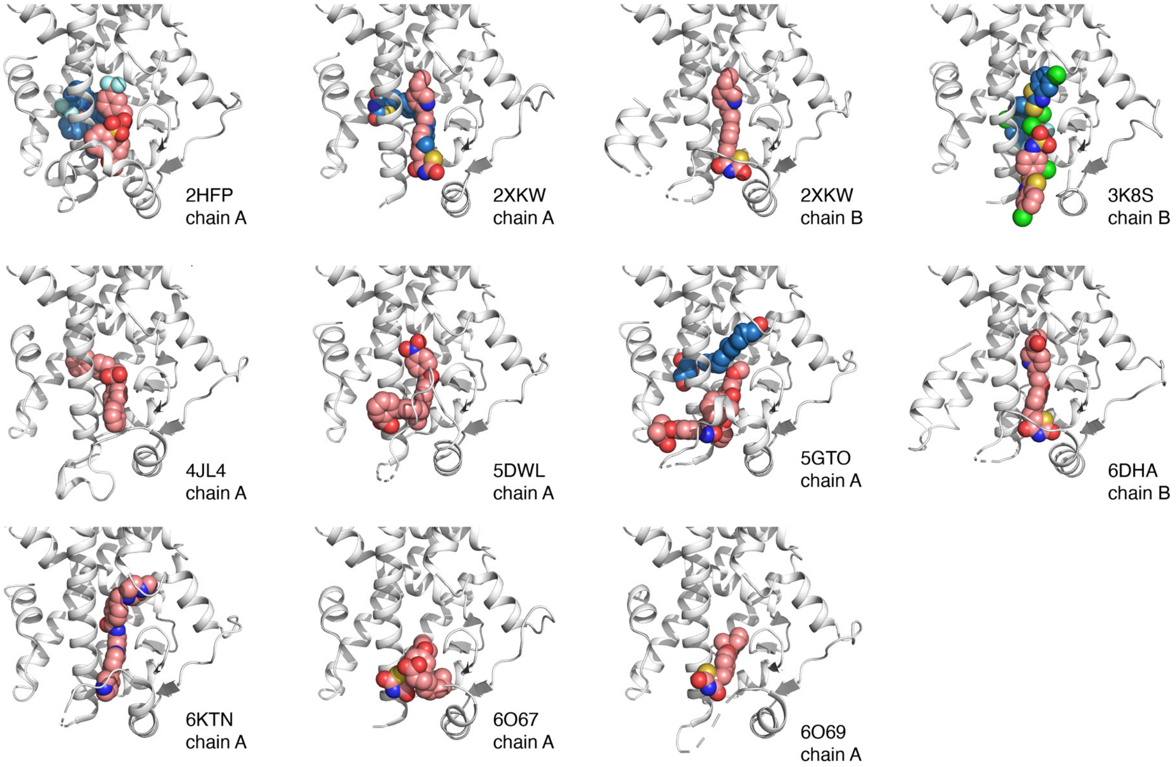

**Fig. S4.** Crystal structures of PPARγ LBD with ligands bound to the orthosteric pocket entrance.

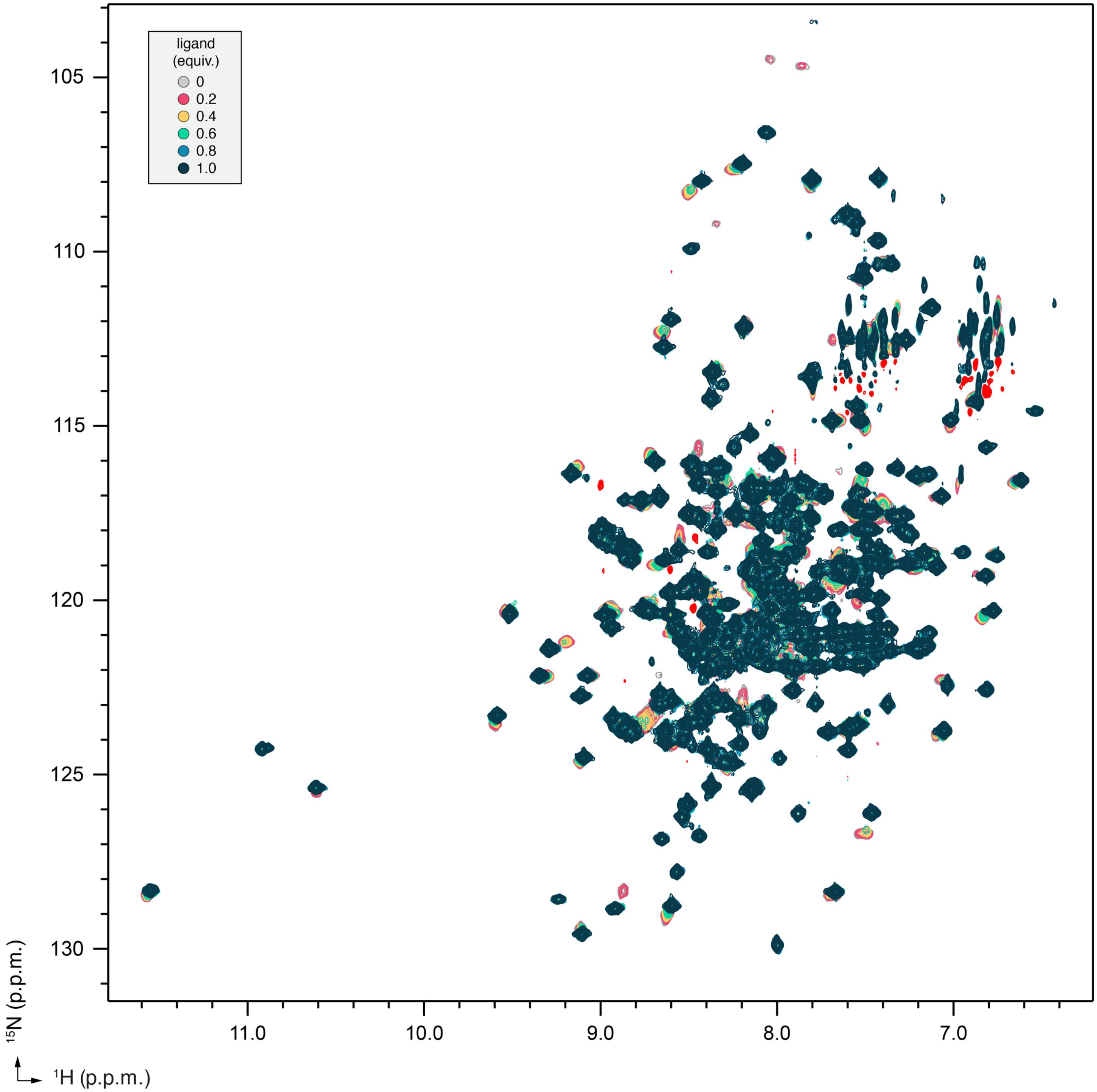

**Fig. S5.** 2D [$^1$H,$^{15}$N]-TROSY-HSQC NMR data of darglitazone titrated into $^{15}$N-PPARγ LBD at the indicated molar equivalents.

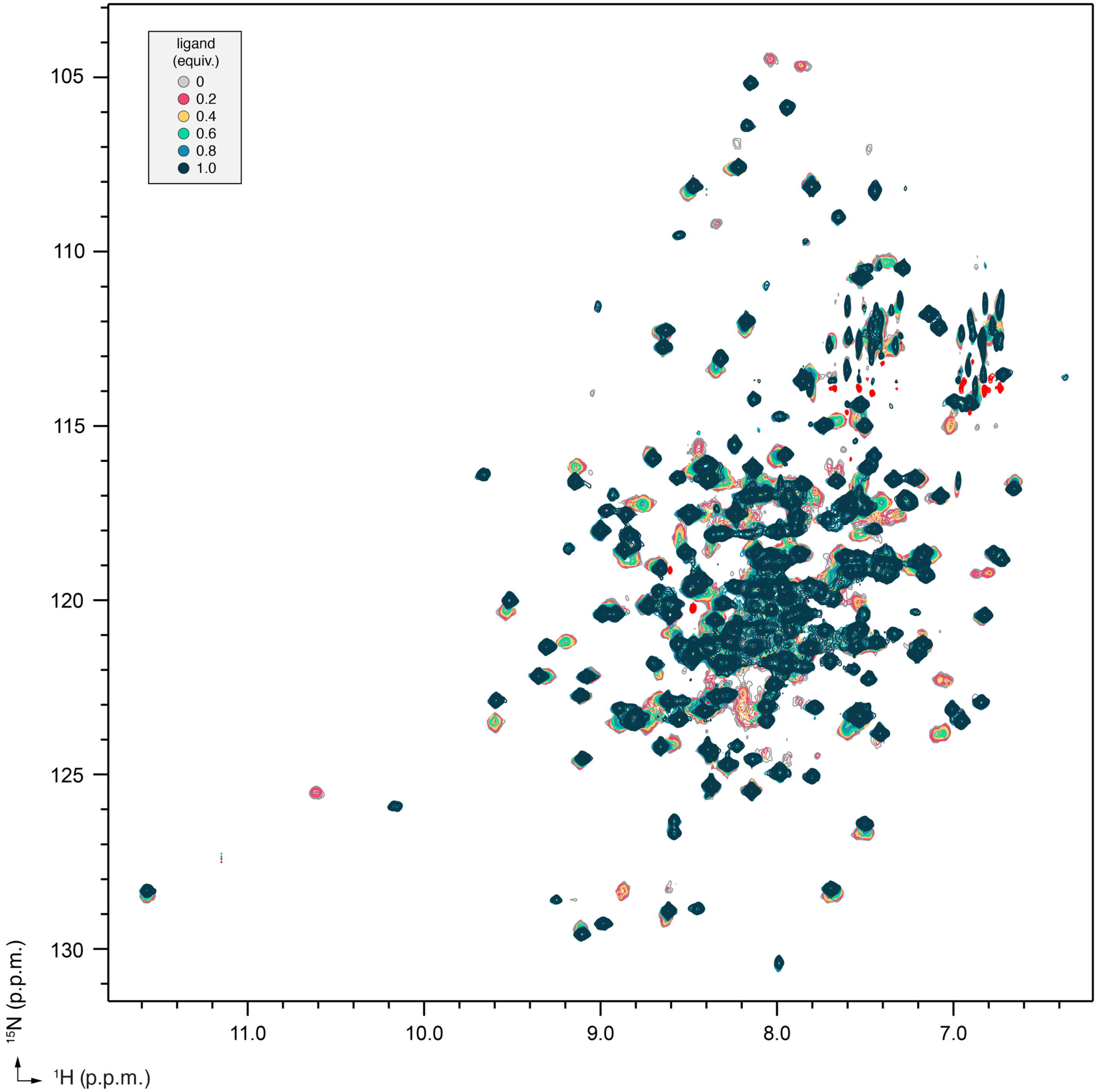

**Fig. S6.** 2D [$^1$H,$^{15}$N]-TROSY-HSQC NMR data of MRL24 titrated into $^{15}$N-PPARγ LBD at the indicated molar equivalents.

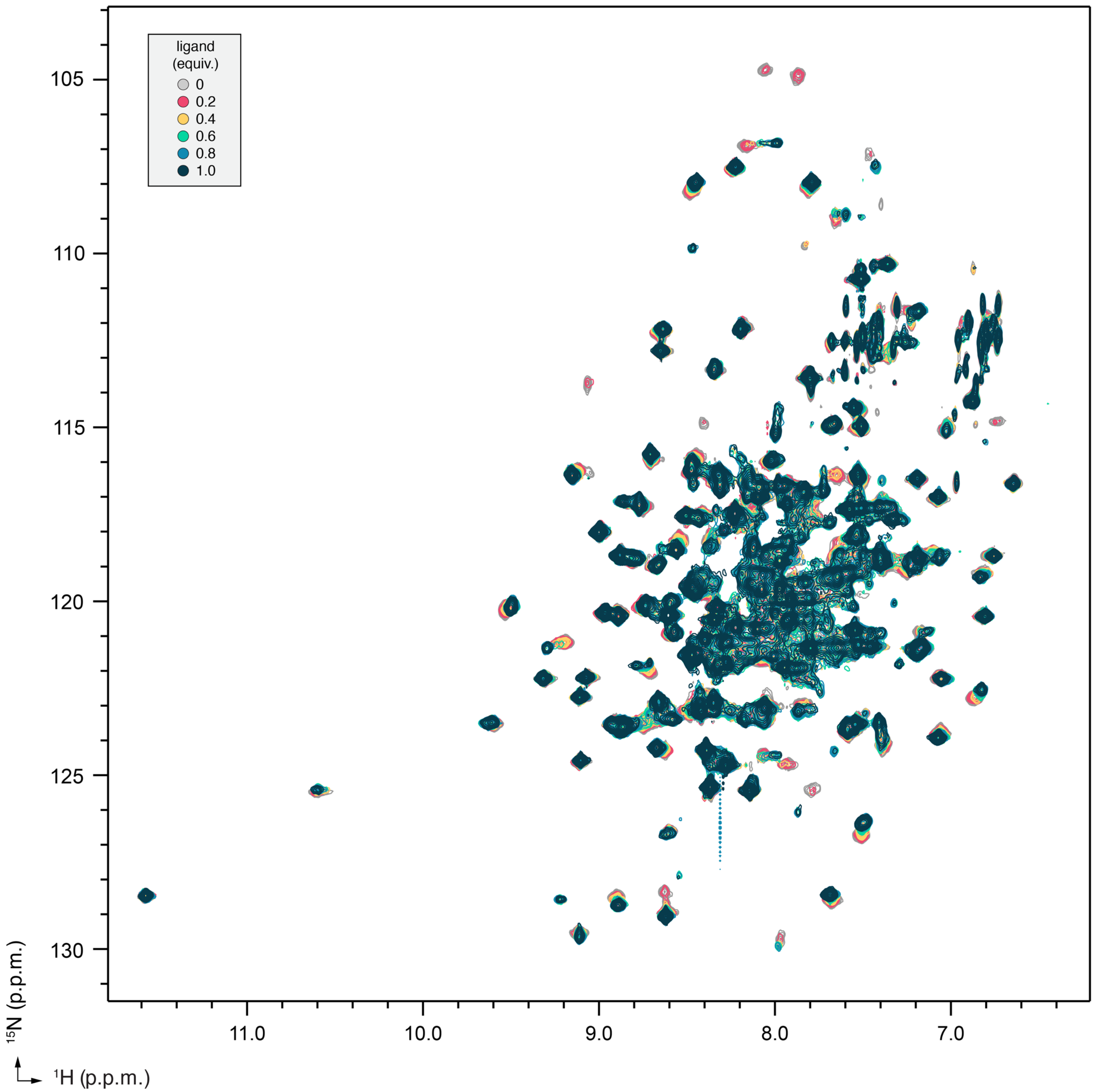

**Fig. S7.** 2D [$^1$H,$^{15}$N]-TROSY-HSQC NMR data of darglitazone titrated into $^{15}$N-[Y473E]-PPARγ LBD at the indicated molar equivalents.

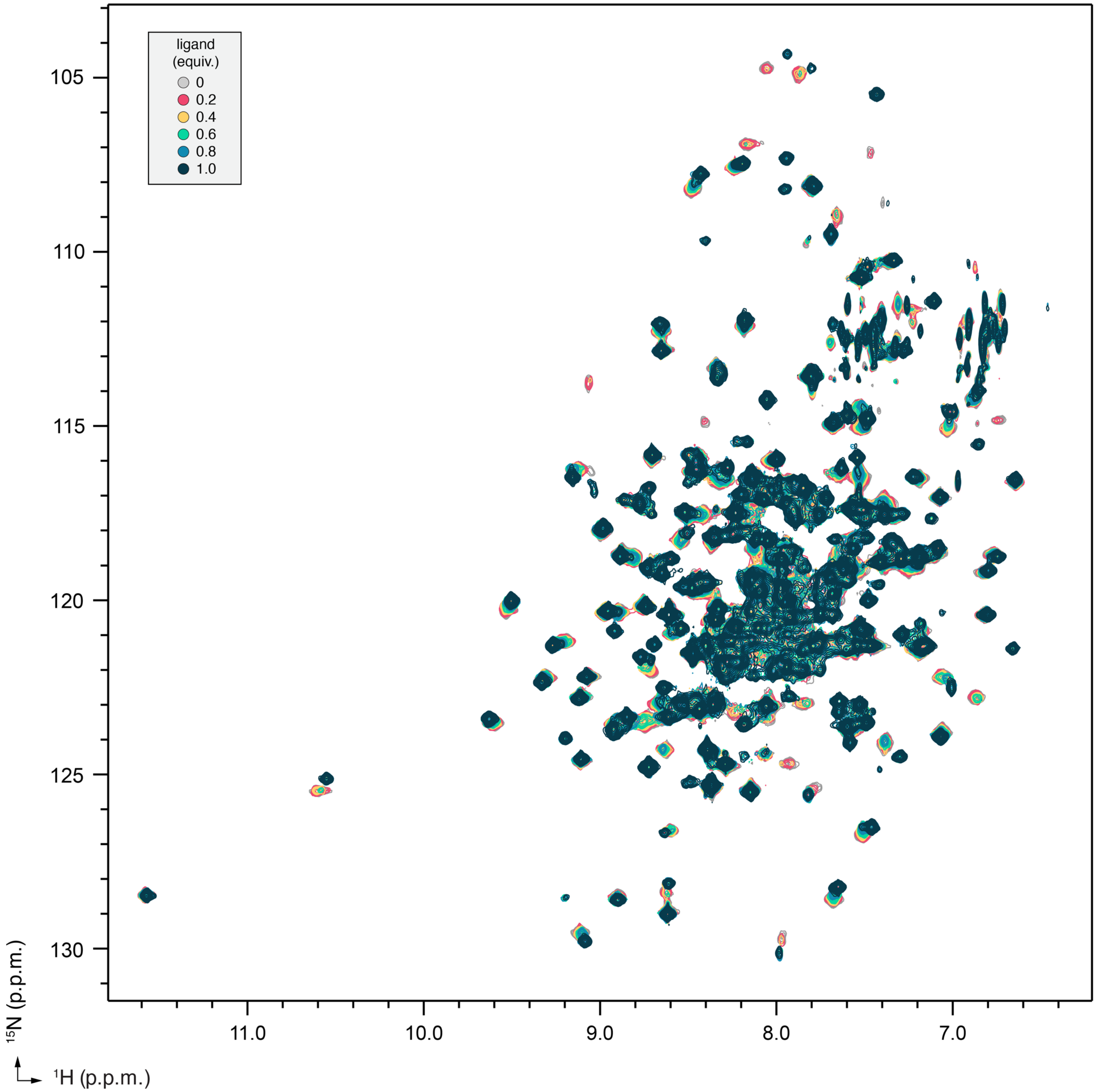

**Fig. S8.** 2D [$^1$H,$^{15}$N]-TROSY-HSQC NMR data of GW1929 titrated into $^{15}$N-[Y473E]-PPARγ LBD at the indicated molar equivalents.

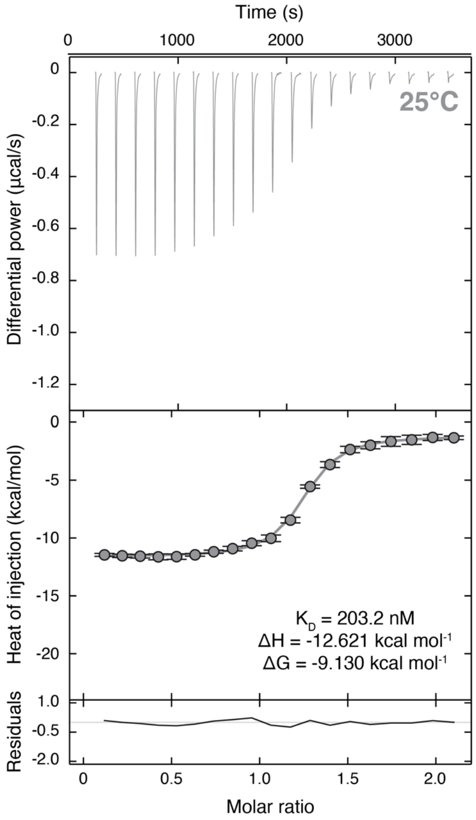

**Fig. S9.** ITC data of GW1929 titrated into [Y473E]-PPARγ LBD at 25°C.

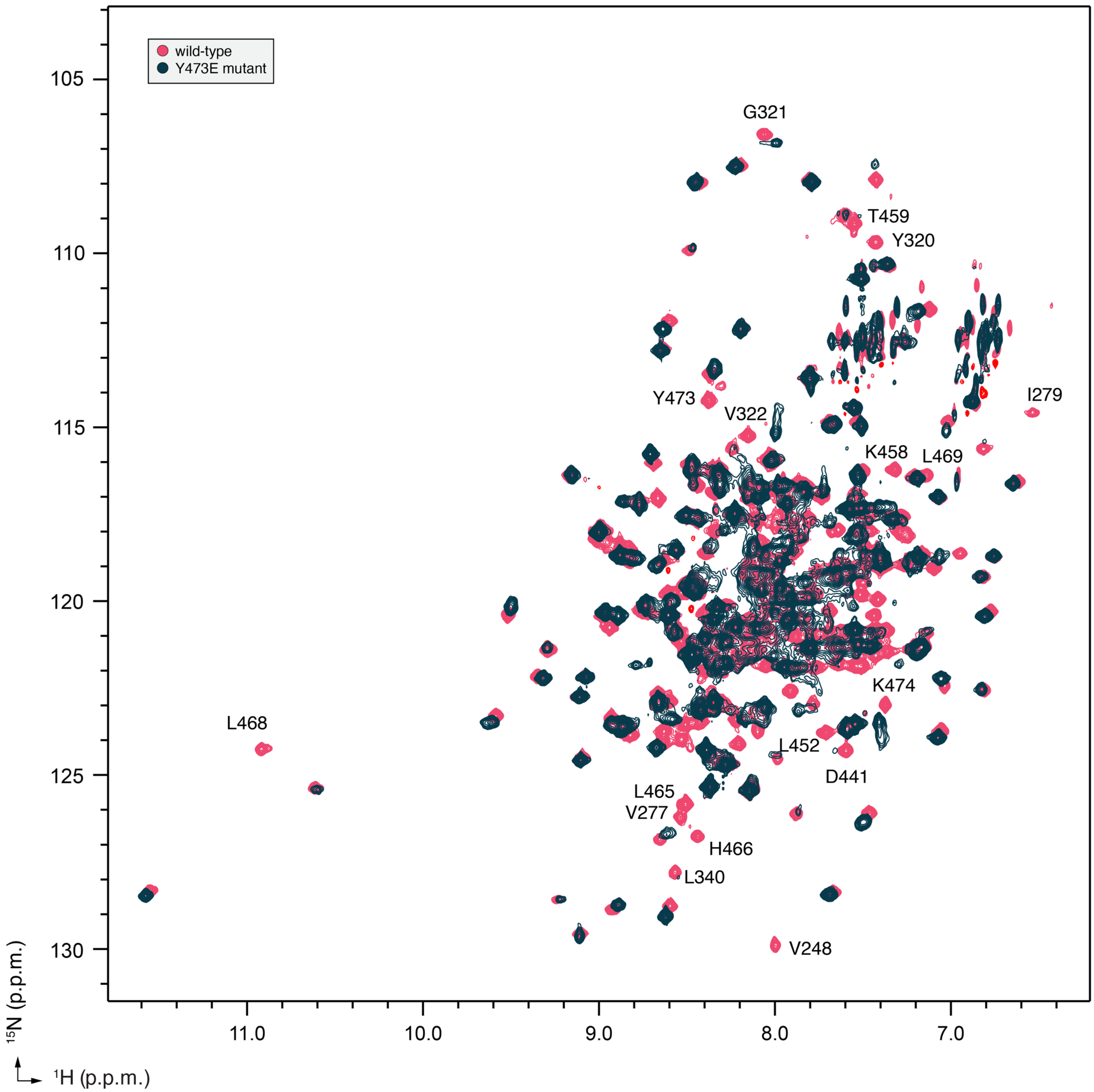

**Fig. S10.** 2D [1H,15N]-TROSY-HSQC NMR data of darglitazone-bound 15N-PPARγ LBD and 15N-[Y473E]-PPARγ LBD.

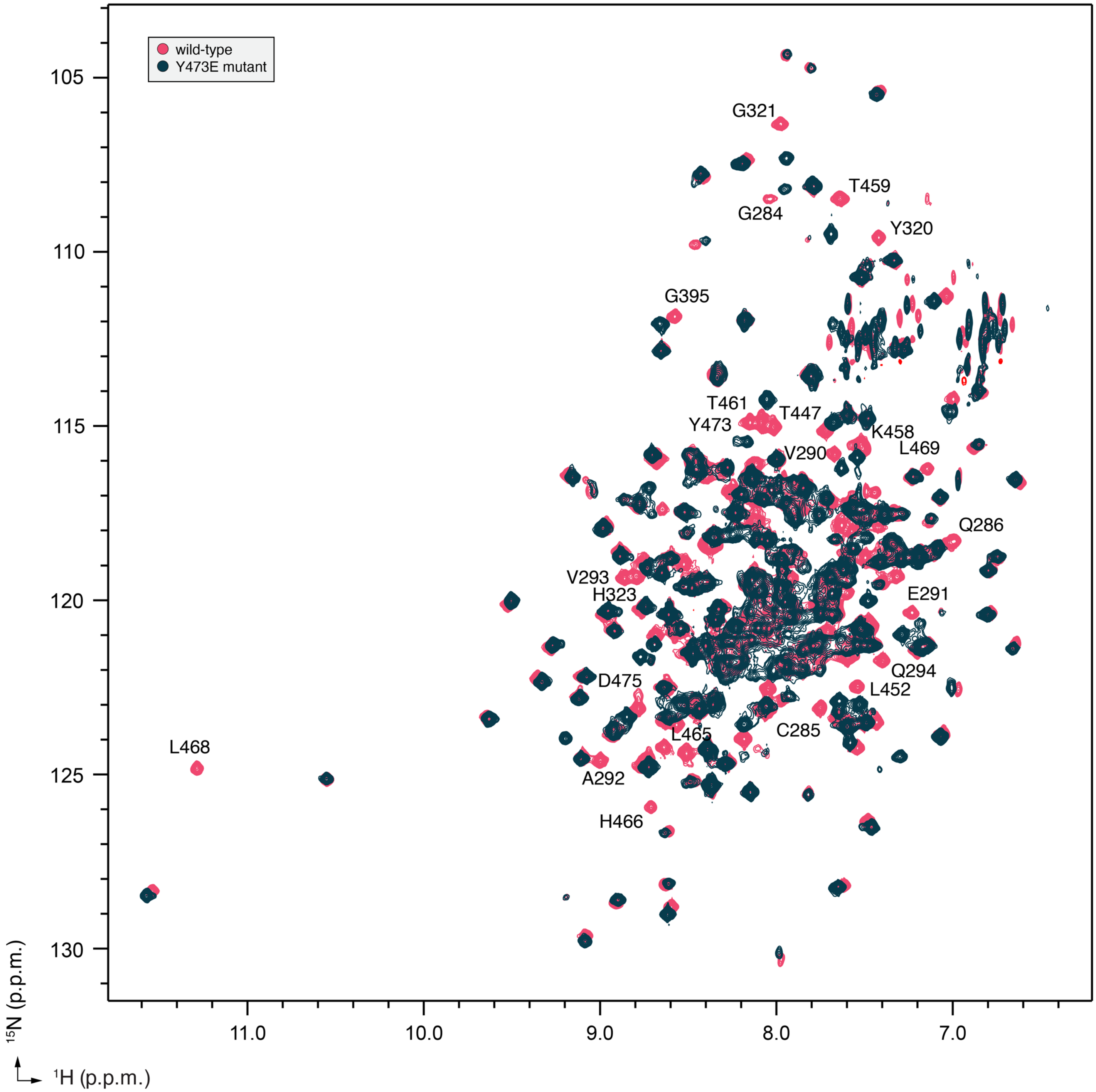

**Fig. S11.** 2D [$^{1}$H,$^{15}$N]-TROSY-HSQC NMR data of GW1929-bound $^{15}$N-PPARγ LBD and $^{15}$N-[Y473E]-PPARγ LBD.

**Table S1. X-ray crystallography data collection and refinement statistics.**

| | PPARγ LBD-Apo Delipidated | PPARγ LBD + GW1929 | [Y473E]-PPARγ LBD | [Y473E]-PPARγ LBD + Darglitazone | [Y473E]-PPARγ LBD + GW1929 |
|---|---|---|---|---|---|
| Data collection | | | | | |
| Space group | C 1 2 1 | C 1 2 1 | C 1 2 1 | C 1 2 1 | C 1 2 1 |
| Cell dimensions | | | | | |
| a, b, c (Å) | 93.04, 62.10, 118.85 | 92.84, 62.13, 119.12 | 92.64, 61.75, 118.78 | 93.07, 61.59, 120.53 | 92.70, 61.62, 118.66 |
| α, β, γ (°) | 90, 102.22, 90 | 90, 102.16, 90 | 90, 101.93, 90 | 90, 102.16, 90 | 90, 102.44, 90 |
| Resolution | 58.08-2.27 (2.35-2.27) | 51.27-2.07 (2.14-2.07) | 33.01-2.3 (2.38-2.3) | 33.41-2.40 (2.49-2.40) | 36.15-2.15 (2.23-2.15) |
| $R_{merge}$ | 0.0367 (0.341) | 0.0511 (1.374) | 0.0163 (0.360) | 0.0245 (0.362) | 0.0134 (0.196) |
| I / σ(I) | 8.43 (1.58) | 13.59 (1.22) | 19.42 (1.94) | 13.59 (2.06) | 22.00 (3.49) |
| CC1/2 in highest shell | 0.862 | 0.727 | 0.877 | 0.774 | 0.934 |
| Completeness (%) | 99.50 (97.78) | 98.75 (97.88) | 97.89 (94.32) | 98.16 (98.76) | 98.19 (98.62) |
| Redundancy | 2.0 (1.9) | 6.4 (6.4) | 2.0 (2.0) | 2.0 (2.0) | 2.0 (2.0) |
| Refinement | | | | | |
| Resolution (Å) | 2.27 | 2.07 | 2.30 | 2.40 | 2.15 |
| No. of unique reflections | 30849 | 40606 | 28880 | 25861 | 35158 |
| $R_{work}/R_{free}$ (%) | 22.3/26.7 | 25.3/28.6 | 22.4/27.5 | 22.2/27.3 | 23.0/26.5 |
| No. of atoms | | | | | |
| Protein | 4122 | 4012 | 4050 | 4075 | 3999 |
| Water | 270 | 335 | 122 | 205 | 90 |
| B-factors | | | | | |
| Protein | 31.34 | 29.54 | 54.96 | 32.78 | 58.40 |
| Ligand | n/a | 53.92 | n/a | 43.47 | 81.00 |
| Water | 31.84 | 31.37 | 50.39 | 31.70 | 54.06 |
| Root mean square deviations | | | | | |
| Bond lengths (Å) | 0.011 | 0.010 | 0.010 | 0.010 | 0.016 |
| Bond angles (°) | 1.18 | 1.12 | 1.08 | 1.18 | 1.30 |
| Ramachandran favored (%) | 96.61 | 96.92 | 97.76 | 96.96 | 96.91 |
| Ramachandran outliers (%) | 0.40 | 0.41 | 0.20 | 0.40 | 0.82 |
| PDB accession code | 6VZO | 6VZL | 6VZN | 6VZM | 7JQG |
| * Values in parentheses indicate highest resolution shell. | | | | | |
